# Efficient, selectable marker free gene targeting in soybean using novel *Ochrobactrum haywardense*-mediated delivery

**DOI:** 10.1101/2021.10.28.466326

**Authors:** Sandeep Kumar, Zhan-Bin Liu, Nathalie Sanyour-Doyel, Brian Lenderts, Andrew Worden, Ajith Anand, Hyeon-Je Cho, Joy Bolar, Charlotte Harris, Lingxia Huang, Aiqiu Xing, Alexandra Richardson

## Abstract

We report robust selectable marker-free gene targeting (GT) system in soybean, one of the most economically important crops. A novel efficient *Ochrobactrum haywardense*-mediated embryonic axis transformation method was used for the delivery of CRISPR-Cas9 components and donor template to regenerate T0 plants in 6-8 weeks after transformation. This approach generated up to 3.4% targeted insertion of the donor sequence into the target locus in T0 plants, with ∼ 90% mutation rate observed at the genomic target site. The GT was demonstrated in two genomic sites using two different donor DNA templates without a need of a selectable marker within the template. High-resolution Southern by Sequencing (SbS) analysis identified T1 plants with precise targeted insertion and without unintended plasmid DNA. Unlike previous low-frequency GT reports in soybean that involved particle bombardment-mediated delivery and extensive selection, the method described here is fast, efficient, reproducible, does not require selectable marker within the donor DNA, and generates non-chimeric plants with heritable GT.

## Introduction

Soybean (*Glycine max*) has become one of the most economically important legume seed crops. In addition to being a major oilseed crop, it also provides more than a quarter of the total protein for the world’s food and animal feed(Graham and Vance, 2003). Due to population growth along with increased societal interest in plant-based protein diets, the demand for soybean is gradually expanding(Ray et al., 2013). However, unlike considerable progress made in increasing the yields of rice, wheat, and maize through the Green Revolution, only limited improvements have been made for soybean(Liu et al., 2020). The yield and other trait improvement in soybean will require complex and precise genetic manipulations to obtain desired plant height, node number, internode length, branch number, disease resistance, and seed size supplemented with high oil and protein contents. Conventional plant breeding has played a major role in making genetic improvement of soybean and will continue to do so in the future. However, genome editing technology could expedite and revolutionize traditional breeding processes and enable markedly improved precision in soybean breeding.

CRISPR (clustered regularly interspaced short palindromic repeats)-Cas9(Jinek et al., 2012) is a robust and versatile genome editing tool to make targeted DNA double-strand breaks (DSBs)(Kim et al., 1996; Epinat et al., 2003; Christian et al., 2010; Gasiunas et al., 2012; Jinek et al., 2012). Genomic DSBs in somatic cells are repaired predominantly by non-homologous end joining (NHEJ), which is an error-prone repair pathway often resulting in small insertions or deletions (indels) at the DSB site. Homologous recombination (HR)(Wood, 1996), in contrast, is a precise repair pathway that requires a DNA repair template with flanking sequences homologous to those flanking the genomic DSB. Unlike NHEJ, HR is a very low frequency process in somatic cells used for plant transformation. Consequently, NHEJ-mediated gene mutations, which can be recovered in 30-100% of regenerated plants, have become routine(Puchta, 2017), whereas precise gene targeting (GT) via HR remains inefficient and challenging. Some monogenic simple traits that depend on endogenous gene mutation can be generated through NHEJ-mediated DNA repair (Li et al., 2016; Li et al., 2016; Liu et al., 2017; Nonaka et al., 2017; Li et al., 2018). Complex yield traits require precise GT via HR to make large modifications in the genome to expedite the breeding process(Shi et al., 2017; Yu et al., 2017).

The majority of genome editing work reported in soybean describes successful NHEJ-mediated targeted mutagenesis(Al Amin et al., 2019; Bao et al., 2019; Campbell et al., 2019; Cheng et al., 2019; Di et al., 2019; Do et al., 2019; Han et al., 2019; Li et al., 2019; Cai et al., 2020; Michno et al., 2020; Zheng et al., 2020). So far, very limited success has been obtained in HR-mediated GT in soybean, with the majority of regenerated plants being chimeric and mutations not inherited to the next generation(Li et al., 2015).

Here we report a novel *O. haywardense*-mediated embryonic axis transformation system for high frequency GT in soybean, where the resultant HR-modified target locus is selectable-marker-free. Using two different donor designs 2-3.4% GT was achieved in two soybean genomic sites. Analyses of T1 progeny confirmed Mendelian inheritance for the majority of GT events, while randomly integrated T-DNA segregated independently of the GT locus resulting in T-DNA-free GT-positive T1 progeny. The method being simple, robust, highly efficient and reproducible, could expedite soybean precision breeding for yield and other traits.

## Results

### Gene Targeting (GT) experiment design

An overview of the GT construct design used in this study is described in Figure 1. The T-DNA construct (Figure 1A and Supplementary Figure 1A) targeting soybean DD38(Cigan et al., 2016) genomic site contained five expression cassettes. The expression cassette nearest the LB encoded a *hygromycin*-resistance gene *(hptII*) donor DNA repair template flanked by homology arms, HR1 and HR2, which were in turn flanked by two Cas9 cut sites matching the genomic target site. The T-DNA construct also contained in order between the RB and *hptII* cassette a single guide RNA expression cassette, a Cas9 expression cassette, a *spectinomycin*-resistance gene (*spcN*), and a *DsRED* color marker cassette. Following T-DNA transfer, transient Cas9 expression results in two DSBs at the Cas9 cut sites next to the two homology arms flanking *hptII*. leading to release of the donor DNA repair template. The T-DNA containing *spcN* as the plant selectable marker, which is randomly inserted in the genome, is used for positive selection and regeneration of transformed cells. In addition, a DSB generated at the genomic target locus, which could then be repaired via HR using the donor template released from T-DNA, resulting in GT. The T-DNA could also be initially randomly integrated into the genome, with Cas9 expression leading to the release of the donor DNA repair template resulting in intragenomic GT as described previously(Barone et al., 2020). The donor DNA repair template in T-DNA construct (Figure 1B and Supplementary Figure 1B) targeting soybean DD51(Cigan et al., 2016) genomic site contained yellow fluorescent protein (*YFP*) gene.

**Figure 1.**
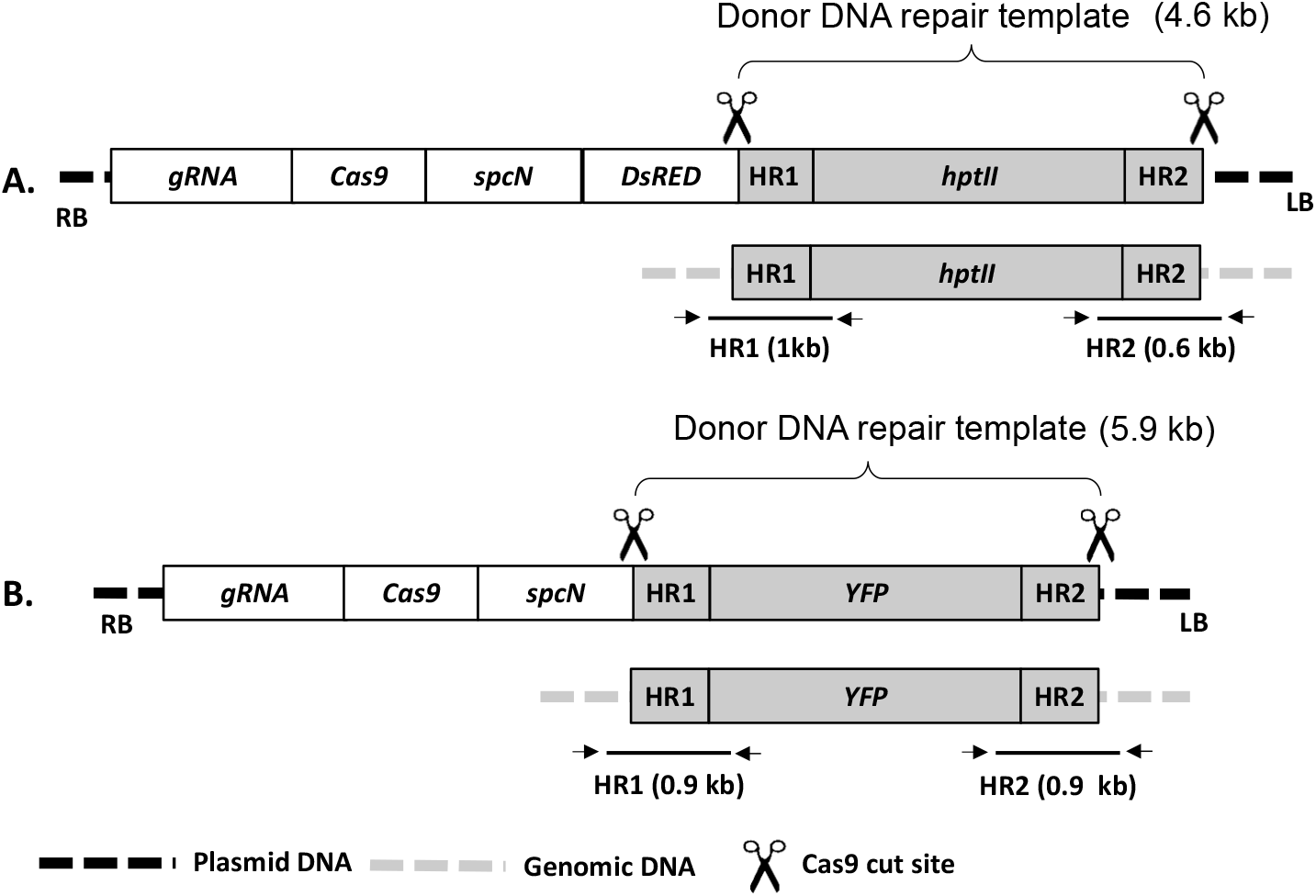
Schematic description of the constructs pDD38 (A) and pDD51 (B) containing donor template for gene targeting at soybeanDD38 and DD51 genomic site, respectively. Both constructs contain guide RNA (gRNA) and Cas9 expression cassettes for genome editing, and *spectinomycin (spcN)* cassette as plant selectable marker. The construct A for DD38 site also contained *DsRED* color marker. Donor DNA repair template contained *Hygromycin (hptII*, A*) or YFP* (B) gene cassette flanked by homology arms (HR1 & HR2) and Cas9 cut sites (shown as scissors) matching DD38 or DD51 genomic site. Following Cas9 and gRNA expression the donor template will be released and used as template for repair at DD38 or DD51 target site for gene targeting (Bottom panel). HR1 and HR2 diagnostic PCRs were done to detect gene targeting as described in Figure 2. RB & LB: Right and left T-DNA border, respectively. The component sizes are not to the scale.

### GT in soybean

A plant transformation construct pDD38 containing constitutively expressed *Cas9* was used for GT at soybean DD38 site (Figure 1A and Supplementary Figure 1A). *O. haywardense*-mediated embryonic axis transformation was carried out to generate stably transformed shoots ∼1.5 cm in size using spectinomycin selection as described in Materials and Methods. In 6-8 weeks after transformation, a total of 466 shoots were regenerated and analyzed for GT and targeted NHEJ-mediated mutation at DD38 site. HR-mediated GT was detected by HR1 and HR2 junction qPCR as depicted in Figure 1A. The junction PCR was designed for quick screening of putative GT positive plants, further analyses for perfect GT was performed in T1 generation. Out of 466 shoots analyzed, HR1 junction-PCR positive product was observed in 48 (10%) shoots, while 43 (9.2%) shoots were found positive for HR2 junction-PCR (Figure 2 and Table 1). Positive PCR products for both HR1 and HR2 junctions were obtained in 16 plants indicating 3.4% putative GT at DD38 site. Targeted insertion mutation analysis of the target site revealed 93.3% of shoots containing mutations in at least one allele, while >7% of the plants were observed with bi-allelic mutations (Table 1).

**Figure 2.**
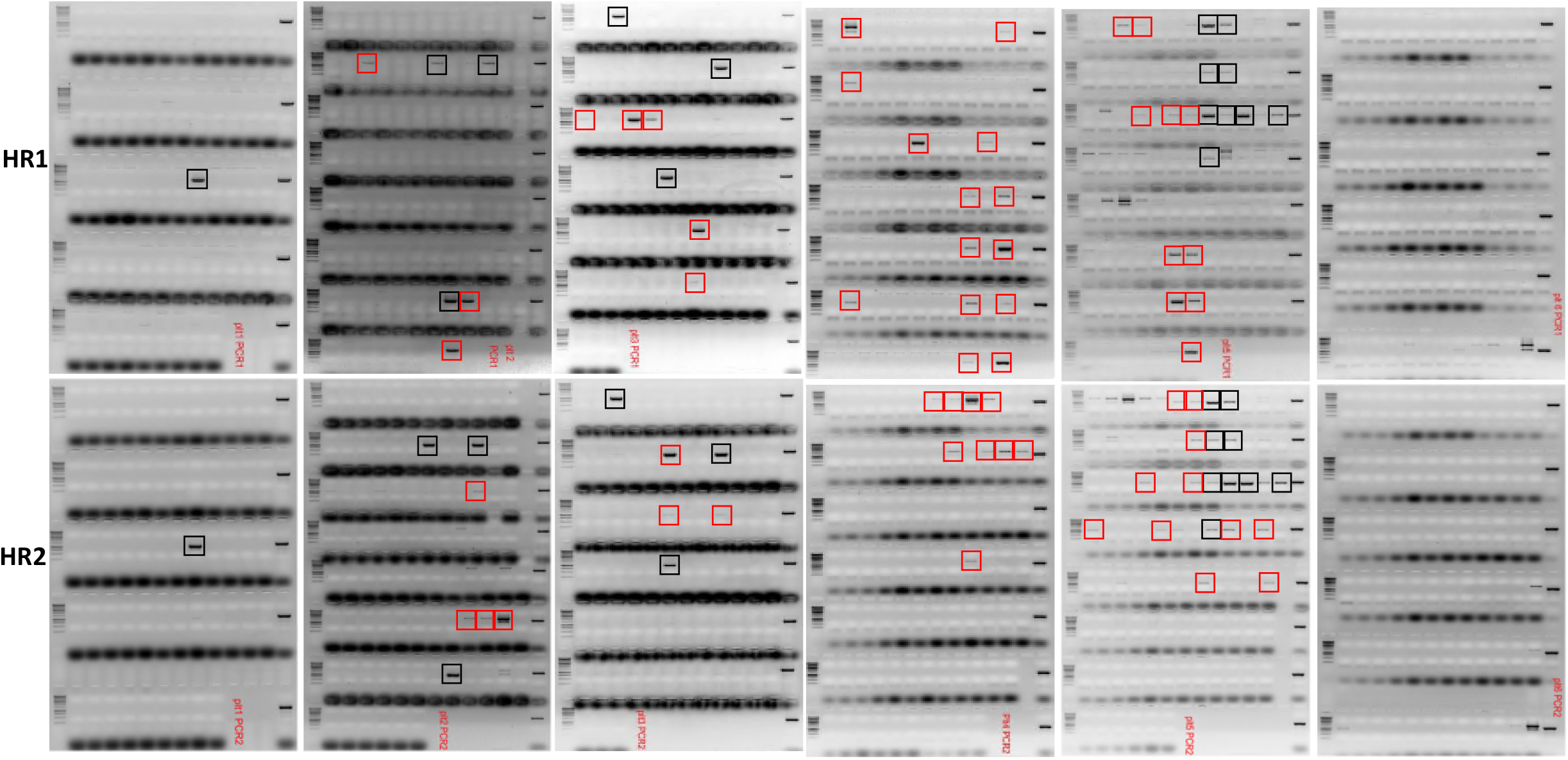
Quick PCR screening to detect gene targeting. HR1 (Upper panel) and HR2 PCR (Lower panel) of leaf samples 466 T0 shoots regenerated for DD38 GT experiment. Positive PCR products of expected sizes obtained both for HR1 and HR2 junctions are shown in black frames. PCR products amplified only for HR1 or HR2 are indicated in red frames. The PCR results are summarized in Table 1.

**Table 1:**
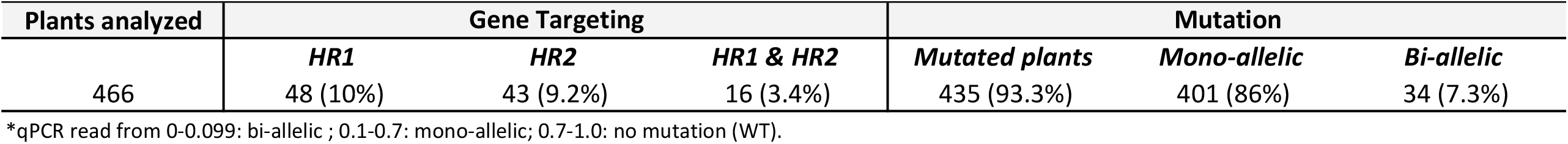
Summary of gene targeting and mutation analyses at soybean DD38 genomic site

After having successfully demonstrated GT at DD38 site, we created another donor construct pDD51 (Figure 1B and Supplementary Figure 1B) targeting DD51 site in soybean genome. The construct pDD51 contained *YFP* cassette as donor repair template while *DsRed* was not included in the T-DNA, the rest of the construct design was similar to pDD38. Using *O. haywardense*-mediated embryonic axis transformation 690 shoots were generated on spectinomycin selection. HR1 and HR2 junction PCR revealed 2.8% and 2% positive plants (Table 2), respectively. Fourteen plants were observed positive for both HR1 and HR2 junctions demonstrating 2% GT at DD51 site. Similar to the DD38 site, high frequency (89%) NHEJ-mediated mutations were detected at the DD51 site, with 9.5% of plants showing bi-allelic mutations.

**Table 2:** Summary of gene targeting and mutation analyses at soybean DD51 genomic site

| Plants analyzed | Gene Targeting |  |  | Mutation |  |  |
| --- | --- | --- | --- | --- | --- | --- |
|  | *HR1* | *HR2* | *HR1 & HR2* | *Mutated plants* | *Mono-allelic* | *Bi-allelic* |
| 690 | 22 (3.2%) | 19 (2.8%) | 14 (2%) | 614 (89%) | 548 (78.5%) | 66 (9.5%) |

\*qPCR read from 0-0.099: bi-allelic ; 0.1-0.7: mono-allelic; 0.7-1.0: no mutation (WT).

### Inheritance and segregation analysis of GT plants

To study inheritance and segregation of GT, four GT positive T0 plants at DD38 site (designated as P1 – P4) were self-pollinated to generate T1 progeny. Approximately 90 T1 plants from each of the 4 GT-positive T0 plants were analyzed for the presence of GT and T-DNA components. To confirm GT and validate integrity of the insertion, HR1/HR2 junction PCR data were supplemented with Southern by Sequencing (SbS(Zastrow-Hayes et al., 2015)) (Table 3). Approximately 75% progeny from 3 GT-positive T0 plants (P2, P3 and P4) were observed positive for both HR1 and HR2 junction PCR, conformed to expected Mendelian inheritance of the GT insertion at DD38 locus. These plants contained 1-2 copies of randomly inserted T-DNA components that segregated 1:2:1 (homo:hemi:null) as expected for a single locus insertion. For two of the four DD38 insertion lines, the number of T1 plants positive for both the HR1 and HR2 junctions and also null for the randomly inserted T-DNA, conformed to the expected Mendelian ratios of ∼25% (16 nulls out of 67 for P3, 22 nulls out of 71 for P4). Only one T-DNA-free GT T1 plant was observed from P2 plant. The T1 progeny from P1 T0 plant did not show expected Mendelian inheritance, only 7 plants (∼7%) were observed GT positive and no T-DNA-free GT T1 plant was obtained (Table 3).

**Table 3:** Inheritance and Segregation of GT in T1 Generation

| *T0 plant* | *Total T1 plants* | **T-DNA** |  |  |  | **Gene targeting** |  |  |
| --- | --- | --- | --- | --- | --- | --- | --- | --- |
|  |  | *Nulls* | *Homo* | *Hemi* | *Copy# (PSB1)* | *HR1/2* | *Positive T1 plants* | *SBS\** |
|  |  |  |  |  |  | *Total* | *Trangene free* |  |
| P1 | 91 | 31 | 21 | 39 | 3 | 7 | 1<sup>φ</sup> | Deletion |
| P2 | 93 | 19 | 16 | 58 | 1 | 71 | 1 | Clean |
| P3 | 86 | 19 | 26 | 41 | 2 | 67 | 16 | Rearrangement |
| P4 | 95 | 26 | 20 | 49 | 2 | 71 | 22 | Clean |

<sup>φ</sup>HR2 only
\*Maximum 5 plants analyzed

Further molecular characterization of T-DNA component-free GT plants was done using SbS, which utilizes in-solution sequence capture coupled with NGS(Zastrow-Hayes et al., 2015). The method is highly sensitive in detection of construct-to-genome and construct-to-construct novel junctions that could have been created during the transformation process. The junction sequence data is then used to detect unintentionally inserted sequences from the transformation plasmid and help identify small T-DNA fragments, truncations of the intended donor DNA for GT, or presence of transformation plasmid backbone sequences. One T1 plant each from P1 and P2 and five T1 plants each from P3 and P4 T0 plants were sampled for SbS (Table 3). The P1 plant although PCR-negative for HR1 was included to confirm sensitivity of SbS. In addition, samples from wild-type negative control and plasmid-spiked positive control were also included.

The sequence coverage graphs of the GT plants and controls were mapped to expected schematic of precise donor insertion at DD38 genomic site (Figure 3 A) and to schematic of the construct used (Figure 3B) to confirm precise donor DNA insertion and absence of unintentional randomly inserted construct sequences. The SbS analyses revealed T-DNA-free precise GT in P2 and four P4 plants A representative result is shown in Figure 3AIII and BIII. One T1 plant from P4 was observed to contain unexpected unique junction at DD38 target site indicating insertion of multiple copies of rearranged donor DNA (Figure 3 A VI). Similarly, all five P3 progeny plants were found to contain rearranged donor DNA at DD38 target site (Figure 3 A V). In addition, these plants also contained small T-DNA and backbone fragments (Figure 3 B V) that were undetected by quick PCR screening conducted in T0 generation. As expected from the HR1 PCR analysis, one P1 progeny plant was observed to contain partial donor DNA insertion lacking majority of intended donor DNA sequence (Figure 3 A IV).

**Figure 3.**
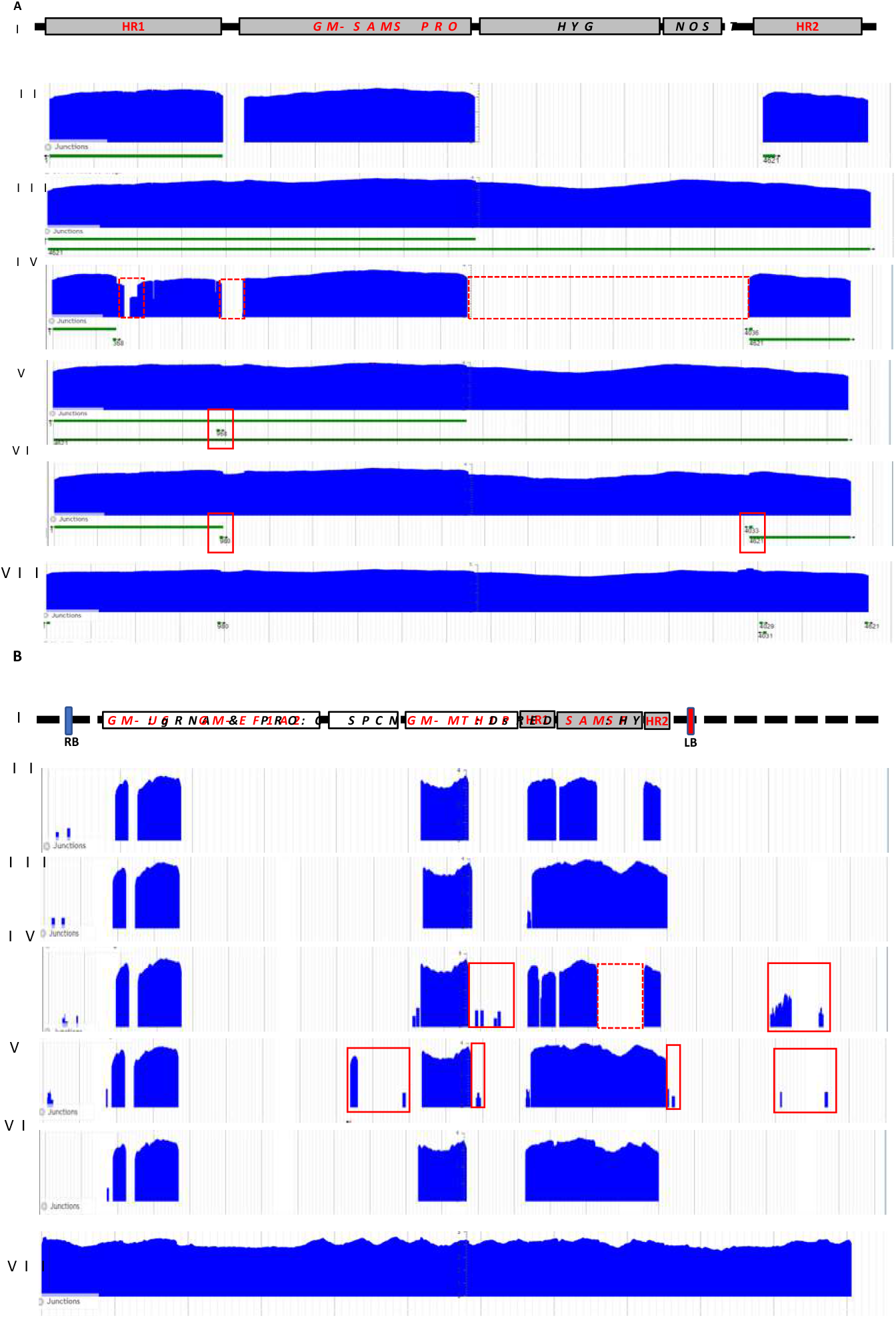
Southern-by-Sequencing (SbS) for high resolution molecular characterization of PCR positive gene targeted T1 soybean plants. **A**. Sequence coverage graphs of targeted plants and controls mapped to expected schematic of precise donor insertion at DD38 genomic site. **I**. Schematic representation of the precisely targeted soybean DD38 site, which is mapped to sequence coverage graph of wild type plant as negative control (**II**), representative precisely targeted P2 T1 plant (**III**), P2 T1 plant observed with imprecise insertion (**IV**) due to missing part of the donor (indicted as dotted red boxes), P3 and P4 T1 plants (**V and VI**, respectively) showing imprecise insertion due to rearrangements at targeted locus (indicted as red boxes), WT plant DNA with spiked in plasmid DNA used for gene targeting as positive SBS control (**VII**). **B**. Sequence coverage graphs of targeted plants and controls mapped to schematic of transgene used. **I**. Schematic representation of plasmid map aligned to sequence coverage graph of plants and controls described in **A. I-VII** above. Red box indicates presence of plasmid DNA, dotted box shows missing part of donor. Red font in the maps indicate soybean genomic sequences

## Discussion

Being a major source of plant proteins and oils, soybean is an economically important food and oilseed crop. The recent societal shift towards the plant-based meat alternatives (Choudhury et al.) has further increased the demand for soybean, making this crop even more important than ever. While Green Revolution has brought a considerable increase in the yields of rice, wheat and maize, only modest yield improvements have been made for soybean(Liu et al., 2020). The unique architecture of soybean plants with leaves, inflorescences, and pods growing at each node, and the determined internode number greatly affect the final yield. Therefore, soybean yield being a multifactorial, complex trait, is likely to require simultaneous and precise modification of several components to positively affect the yield. Modern precision breeding tools, including genome editing, present an opportunity for such targeted breeding approach. In addition, genome editing would be extremely useful in making pathway engineering needed to develop soybean varieties with enhanced protein and oil contents(Aili Bao, 2020; Subedi et al., 2020).

CRISPR-Cas9 due to flexibility and ease of the design has become an important genome engineering tool for creating targeted DSBs required for precision breeding applications(Wiedenheft et al., 2011; Jinek et al., 2012; Cong et al., 2013; Xing et al., 2014). DSB at desired target site triggers induction of a DNA repair pathway. In somatic cells NHEJ remains a dominant pathway, which mainly involves ligation of unrelated sequences or sequences with micro-homologies (Salomon and Puchta, 1998; Puchta, 1999; Orel et al., 2003). Therefore, CRISPR-Cas9-medaited targeted mutagenesis, which is based on the NHEJ repair, is now becoming routine in plants that are amenable for transformation including soybean(Al Amin et al., 2019; Bao et al., 2019; Campbell et al., 2019; Cheng et al., 2019; Di et al., 2019; Do et al., 2019; Han et al., 2019; Li et al., 2019; Cai et al., 2020; Michno et al., 2020; Zheng et al., 2020). In contrast, GT for targeted insertion or allele replacement remains a huge challenge mainly due to low-frequency HR in somatic cells coupled with complexity of making externally supplied donor template accessible at DSB site(Mao et al., 2019; Rozov et al., 2019). The complex genetic changes needed to improve soybean yield, oil and protein traits are likely to require precise insertions and replacements of DNA sequences, which can only be accomplished by HR-mediated GT. The challenge of inefficient GT process is further confounded by the need of a selectable marker to recover, enrich and regenerate plants from rare GT cell(s), while the presence of the selectable marker in the final GT plants is undesirable for commercial deployment of the technology. Therefore, development of a GT system which does not have a selectable marker in the final plant is highly desirable for application of GT for commercial product development.

Here we report novel *O. haywardense* -mediated delivery of CRISPR-Cas9 and donor DNA template to obtain high frequency GT in soybean. The delivery method utilized embryonic axis derived from soybean seed to generate stably transformed shoots (Cho et al. Manuscript submitted). First, we demonstrated efficient GT at soybean DD38 genomic site. While up to 10% targeted insertion was observed by HR1 or HR2 junction PCR, 3.4% putative true GT was obtained as indicated by positive PCR for both HR1 and HR2 junctions. Second, to establish reproducibility of our GT method we targeted another genomic site (DD51) using a different construct design. Similar to DD38 site, we successfully generated 2% GT at DD51 site indicating that GT method described in this report is reliable and can be applicable for different genomic sites. High frequency (∼90%) targeted mutagenesis both at DD38 and DD51 sites further confirm that our CRISPR-Cas9 genome editing is robust and efficient.

While the donor DNA template contained *hptII* or *YFP* gene cassette, they were not used for selection; instead, *spcN* located on the T-DNA outside the donor DNA sequence was used to identify plants with randomly inserted T-DNA. Such construct design enables the donor DNA to contain only the trait gene(s) and the GT at genomic target site to be free from selectable marker. The randomly inserted T-DNA containing *spcN* cassette can be segregated out in next generation to obtain clean GT plants without unintended T-DNA sequences. The unique donor construct design with Cas9 cut sites flanking donor template(Peterson et al., 2021) enabled such indirect selection using randomly inserted *spcN* selection cassette while GT at genomic target site remains selection free. Recently, we have applied similar construct design for an intra-genomic GT in maize, however, it utilized a heat-shock inducible expression of Cas9(Barone et al., 2020). The GT work on soybean described in this paper is based on the constitutive expression of Cas9. While we assume that selective regeneration advantage for the Cas9-expressing cells with donor released from T-DNA provided preferential enrichment for GT events, intra-genomic GT with donor released from the T-DNA randomly integrated into the plant genome cannot be ruled out. Therefore, similar to our maize intra-genomic GT(Barone et al., 2020), inducible Cas9 expression coupled with selectable marker activation would be an attractive approach to further enhance GT in soybean.

Lack of fast, efficient and reproducible transformation method has been a key bottleneck for soybean genome editing(Altpeter et al., 2016). The common transformation method used to transform soybean utilizes *Agrobacterium*-mediated DNA delivery to cotyledonary-node explants(Paz et al., 2006), which is not only inefficient (∼4% transformation frequency) but often leads to chimerism and non-transgenic escape plants, making the transformation process labor-intensive and expensive(Zheng et al., 2020). The current status of soybean transformation probably explains the lack of a viable GT system for commercial product development. The only previously published report on GT in soybean utilized particle bombardment-mediated delivery, showing GT in callus stage with majority being chimeric and not inherited to next generation(Li et al., 2015). In contrast, novel *O. haywardense* H1-8 mediated delivery used here is fast, highly efficient (∼35% transformation frequency), reproducible, and generates non-chimeric plants (Cho et al. Manuscript submitted). The transformation method coupled with the construct design described in this report was instrumental for successful demonstration of efficient, heritable, and selectable marker free GT system paving the way for next-generation precision breeding in soybean.

After demonstrating a successful GT in T0 plants, the next step was to validate that our GT method produces non-chimeric plants with stable and precise insertion, while unintentionally randomly inserted T-DNA sequences are segregated away to generate transgene-free GT plants in T1 generation. Among self-pollinated T1 progeny planted from 4 GT-positive T0 plants, Mendelian inheritance of GT at DD38 locus and randomly inserted T-DNA was observed for progeny from 3 T0 plants. Expected 25% T-DNA-free GT positive T1 plants were obtained from two T0 GT plants. Only one T-DNA-free GT plant was observed from the third plant indicating genetic linkage between DD38 target site and T-DNA insertion site in this line. The T1 progeny from one plant showed chimeric GT and no T-DNA-free GT T1 plant was obtained in this line. The next goal was to confirm the fidelity of PCR-positive GT call and ensure selected GT T1 plants are free from any unintentionally inserted T-DNA sequences. We employed high-resolution SbS(Zastrow-Hayes et al., 2015) to further characterize selected transgene-free GT positive T1 plants. Compared to the whole-genome sequencing, SbS utilizes sequence capture technology, which reduces sequence analysis complexity by enriching the target sequences of interest, in turn decreasing the amount of sequence data generated by NGS technology. Bioinformatic analysis of the targeted sequencing identifies novel plasmid-to-genome or plasmid-to-plasmid junctions providing comprehensive information about the number of unique insertion loci, potential rearrangements of the inserted DNA, and the presence or absence of plasmid backbone sequences. SBS analyses revealed random T-DNA-free precise GT in T1 plants from two T0 lines. Some T1 plants from these lines were also observed with insertion of multiple copies of rearranged donor DNA. All T1 plants screened for one T0 line had rearranged donor DNA at DD38 target site, in addition to containing small T-DNA and backbone fragments that were undetected by quick PCR screening in T0 generation. More extensive diagnostic PCR in T0 could have detected these imperfect GT or T-DNA fragment-containing events.

In summary, using a novel *O. haywardense* H1-8 mediated delivery we have developed efficient and reliable GT system in a major dicot crop plant. This work also provides general analytical guidelines for T0 to T1 generations post transformation for selecting plants with precise GT and void of any unintentionally inserted plasmid DNA sequences. Given gene targeting without a selectable marker gene or any transgene in the final plant is required for non-transgenic applications of genome editing technology, successful first demonstration of the system design for a selectable marker-free GT in this report is a major step towards commercial application of CRISPR-Cas9 genome editing in soybean precision breeding.

## Methods

### Plasmids and reagents used for plant transformation

Standard cloning methods were used to construct DNA plasmids used in this study. All plasmids were quality controlled through deep sequencing and 100% identity was confirmed. Supplementary Table 1 and Supplementary Figure 2 provide the details of expression and other elements used in different constructs described in Figure 1 and Supplementary Figure 1.

### Soybean transformation

Corteva Agriscience elite soybean variety 93Y21 was used. Mature dry seeds were surface-sterilized for 16 hours using chlorine gas as described by Di et al.(Di et al., 1996). Disinfected seeds were imbibed on solid agar medium containing 5 g/L sucrose and 6 g/L agar for 6-8 hrs and then the seeds were soaked in distilled sterile water for overnight at room temperature in the dark. Intact embryonic axes were isolated using a scalpel blade.

All *O. haywardense* H1-8 -mediated soybean transformation was carried out prior to 2020 as described by Cho et al. (Cho et al. Manuscript submitted). *O. haywardense* H1-8 lines containing the vectors listed in Figure 1 and Supplementary Figure 1 were used for transformation. *O. haywardense* H1-8 suspension (OD 0.5 at 600 nm) in infection medium composed of 1/10X Gamborg B5 basal medium, 30 g/L sucrose, 20 mM MES, 0.25 mg/L GA3, 1.67 mg/L BAP, 200 μM acetosyringone and 1 mM dithiothreitol in pH 5.4 was added to about 200-300 EAs on a 25 × 100 mm deep petri dish. The plates were sealed with parafilm, then sonicated for 30 seconds. After sonication, EAs were incubated 2 hrs at room temperature. After inoculation, about 200-300 EAs were transferred to a single layer of autoclaved sterile filter paper (Cat No. 28320-020, VWR) in 25 × 100 mm petri dish. The plates were sealed with Micropore tape (1530-0, 3M, St. Paul, MN, USA) and incubated under dim light (1-2 μE/m^2^/s), cool white fluorescent lamps for 16 hours at 21°C for 3 days. After co-cultivation, the base of each embryonic axis was embedded in shoot induction medium (R7100, PhytoTech Labs) containing 30 g/L sucrose, 6 g/L agar, 25 mg/L spectinomycin as a selectable agent and 250 mg/L cefotaxime (GoldBio, ST Louis, MO, USA) in pH5.7. Shoot induction was carried out in growth room at 26°C with a photoperiod of 16 hours and a light intensity of 60 - 100 μE/m^2^/s.

After 4-6 weeks in selection medium, the spectinomycin-resistant shoots were cut and transferred to ½ strength MS rooting medium (M404, PhytoTech Labs) containing 15 g/L sucrose, agar 6 g/L, 10 mg/L spectinomycin and 250 mg/L cefotaxime for further shoot and root elongations. Transformation efficiency was calculated based on the number of positive transgenic soybean T0 plants divided by the total number of EAs. Transgenic soybean plantlets were transferred to moistened Berger BM2 soil (Berger, Saint-Modeste, QC, Canada), and hardened plantlets were potted in 2-gallon pots containing moistened SunGro 702 and grown to maturity for harvest in a greenhouse.

### Molecular analysis

Plants were sampled by leaf punching at 7 days after transplantation for quantitative PCR (qPCR) to confirm the presence of the selectable marker (*spcN*), CRISPR-Cas9 (PSN1), the left (PSB1) and right border (PSA2) regions. The *spcN*, PSN1, PSB1, and PSA2 regions for qPCR are shown in T-DNA maps in Supplementary Figure 1 and PCR conditions are given in Wu et al. (2014)(Wu et al., 2014). The primers and probes used for qPCR assays are described in Supplementary Table 2. DNA was extracted from the 7 mm leaf disk with sbeadex™ chemistry (LGC Biosearch, Middlesex, United Kingdom) as per vendor recommendations and resulting DNA was analyzed for quality and quantity via DropSense (Unchained Labs, Pleasanton, CA). PCR to detect mutation and HR junction PCR have been described previously(Barone et al., 2020). Mutation frequencies at genomic target sites were analyzed by qPCR using primer pair and probes given in Supplementary Table 3. HR Junctions were analyzed by using HR spanning primers (F1 and R1) described in in Supplementary Table 3.

Southern-by-sequencing data were generated according to Zastrow-Hayes et al.(Zastrow-Hayes et al., 2015). Briefly, genomic DNA was isolated from 4-mm leaf punches using the Sbeadex Maxi Plant kit (LGC Genomics) then randomly sheared to an average size of ∼400bps with a Covaris E210 ultrasonicator (Covaris). Indexed Illumina genomic libraries were generated with the KAPA HTP Library Preparation Kit (Roche). Adapter-ligated DNA fragments were subsequently PCR-amplified for 8 cycles, pooled in equal molar ratios then normalized in nuclease-free water to 5ng/μl. Following library construction, pooling and normalization, target enrichment was performed using a modified Roche Nimblegen double capture protocol(Zastrow-Hayes et al., 2015), with biotinylated DNA probes designed from the entire plasmid sequences. Enriched DNA fragments were sequenced on an Illumina NextSeq sequencer, according to the manufacturer’s instructions. The resulting DNA sequences were quality-trimmed and aligned to the plasmid sequence for coverage analysis and characterization of the event. Positive SbS controls were derived from wild-type genomic DNA combined with spiked-in plasmid DNA for Illumina genomic library construction and enrichment.

## Authors’ contributions

S.K., Z.B.L., and A.A designed the experiment. N.S.D, J.B. and H.J.C. performed the soybean transformation. B.L., A.W. and A.R. designed and performed DNA analysis. S.K. led the project and wrote the manuscript. All the authors reviewed the manuscript.

## Acknowledgements

We thank Lijuan Wang and Super Vector team for vector construction, Shujun Chang and Sam Ellis for soybean transformation support, Joel Van Wyk, and Becca Vickroy for greenhouse care, and Marv Taylor and Balaji Boovaraghan and the PCR Analysis and Characterization team for high throughput PCR support. The authors also thank Scott Betts, Maria Fedorova, and Sendil Devadas for critical review of the manuscript. We acknowledge Clara Alarcon, Todd Jones and Doane Chilcoat for providing resources and support for this work.

## Data availability

The data supporting the findings of this study are available within the paper and its supplementary information files.

